# Higher-order thalamic implication in the processing of bilateral sensory events

**DOI:** 10.1101/2020.05.01.073098

**Authors:** Carlos Castejon, Angel Nuñez

## Abstract

In the rodent whisker system, it is well assumed that VPM and POm encode stimulations of the contralateral whisker pad. However, during tactile exploration whiskers are usually stimulated bilaterally. Accordingly, the integration of tactile information from the two sides of the body seems to be fundamental in the processing of these events. Here, to investigate whether POm could be able to codify these bilateral dynamics, whisker-evoked responses in this thalamic nucleus were examined by *in vivo* extracellular recordings in anesthetized rats using contralateral and ipsilateral stimuli. Strikingly, we found that POm is also able to respond to tactile stimulation of ipsilateral whiskers. Our findings reveal the implication of POm in the representation of bilateral tactile events by integrating simultaneous signals arising from both whisker pads and demonstrate the implication of the higher-order sensory thalamus in the encoding of bilateral sensory events. This can have important implications in bilateral perceptual function.

## Introduction

Rodents have an array of whiskers on each side of the face. In these animals, whisker information is processed by two main parallel ascending pathways towards the cortex (Diamond et al., 1992; Veinante et al., 2000; Wimmer et al. 2010; Casas-Torremocha et al. 2019). The lemniscal pathway includes the ventral posteromedial thalamic nucleus (VPM). The paralemniscal pathway includes the posteromedial thalamic complex (POm). It is well assumed that VPM and POm encode stimulations of the contralateral whisker pad. However, during tactile exploration whiskers are usually stimulated bilaterally. Accordingly, the integration of tactile information from the two sides of the body seems to be fundamental in the processing of these events in bilateral perceptual function. The somatosensory cortical implication in the processing of bilateral stimuli has been much more studied (Armstrong-James and George 1988; Shuler et al. 2001; Debowska et al. 2011). However, the implication of the thalamus in these tactile interactions remains unknown.

Although the function of VPM has been largely studied, the function of POm is still unclear. Here, to investigate whether POm could be able to codify these bilateral dynamics, whisker-evoked responses in this nucleus were examined by *in vivo* extracellular recordings in anesthetized rats using contralateral and ipsilateral stimuli. We found that POm is also able to respond to tactile stimulation of ipsilateral whiskers. Moreover, our results showed that sensory information from both whisker pads is integrated by POm. The integration of contra- and ipsilateral whisker inputs has been described previously in the barrel cortex (Shuler et al. 2002). However, the implication of the thalamus in this integration had not been described before. Here, our observations show that POm mediates bilateral sensory processing and demonstrate a thalamic implication in bilateral tactile perception.

## Results

### 1. POm responses to ipsilateral whisker stimulation

First, whisker-evoked responses in POm were examined using contralateral stimuli. In agreement with previous findings (Chiaia et al. 1991; Ahissar et al. 2000), we found robust whisker-evoked responses with very short latencies (mean response onset latency: 11.78 ± 0.71 ms, n = 85; Fig. 1C). These short latency spikes are consistent with direct ascending driving inputs from the trigeminal nuclei. POm responses lasted the duration of the stimuli (Castejon et al. 2016; Fig. 1C). Then, ipsilateral stimulation was applied and unexpectedly, we found that POm was also able to respond to tactile stimulation of ipsilateral whiskers. Ipsilateral responses were less strong in magnitude than contralateral responses (−35 %; mean contralateral response magnitude: 7.93 ± 0.53 spikes/stimulus; mean ipsilateral response magnitude: 5.12 ± 0.37 spikes/stimulus; p<0.001, Wilcoxon matched-pairs test, n = 85; Fig. 1F) and longer in latency (ipsilateral mean response onset latency: 22.40 ± 1.3 ms; Fig.1C, D). These findings were consistent across all animals (n = 8). Importantly, we found that POm responses also lasted the duration of the ipsilateral stimulus (Fig. 1C, F).

**Fig. 1.**
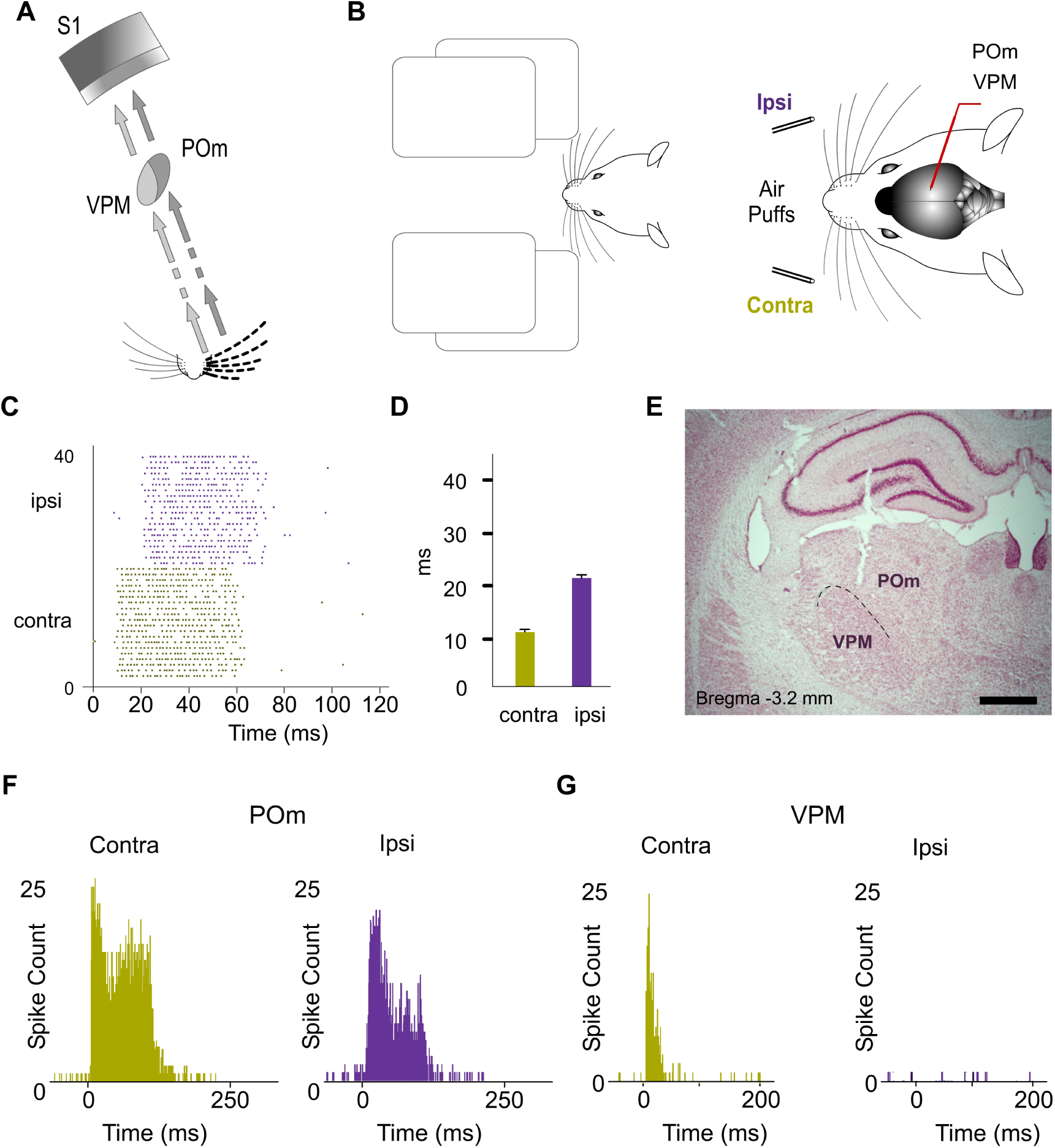
POm responses to tactile stimulation of ipsilateral whiskers. (A) Schematic illustration of the lemniscal and paralemniscal pathways. (B) Usually tactile sensory events are characterized by bilateral whisker patterns. Schematic drawing displaying the sensory stimulation via patterns of contra- and ipsilateral deflections to investigate whether the thalamus could be implicated in the processing of these bilateral events. Recordings were made in the POm and VPM nuclei. (C) Raster plots showing POm responses evoked by contra- and ipsilateral stimuli (stimulus duration 50 ms; 20 trials shown for each stimulus). Note that response onset latency was longer for ipsilateral than contralateral whisker stimulation. (D) Mean response onset latencies of contra- and ipsilateral responses are shown (ipsilateral, n = 85; contralateral, n = 85). (E) Nissl stained coronal section displaying the location of the recording site in the dorsolateral part of POm and the track left by the electrode. Bregma anteroposterior level is indicated. Scale bar, 1 mm. (F) PSTHs showing POm responses evoked by contra- and ipsilateral stimulation (stimulus duration 100 ms; 30 trials shown for each stimulus). Note that although the ipsilateral response was less strong in magnitude, the response remained robust and also lasted the duration of the stimulus. (G) PSTHs showing VPM responses evoked by contra- and ipsilateral stimulation (30 trials shown for each stimulus). VPM did not respond to ipsilateral stimuli

To complement the analyses described above, and as a control, we also characterized VPM thalamic responses delivering the same ipsilateral multiwhisker activation in 6 rats. In contrast to POm, VPM did not respond to ipsilateral stimuli (Fig. 1G). Since the integration of tactile information from the two sides of the body is fundamental in bilateral perception, our results suggest a different implication of these thalamic nuclei in this function.

### 2. POm integration of bilateral inputs

Finally, since bilateral sensory events usually occur simultaneously producing the overlapping of contra- and ipsilateral inputs, the next question to investigate was whether POm could be able to codify this bilateral interaction. It would require the precise integration of contralateral and ipsilateral sensory inputs. To examine the implication of POm in this computation, we applied bilateral sensory stimulation (Fig. 2). We measured the responses of this nucleus to simultaneous contra- and ipsilateral inputs and found that POm integrates tactile events from both sides increasing its response magnitude. Contralateral POm responses exhibited significant increases in firing rate when contra- and ipsilateral whiskers were activated concurrently (Mean response magnitude variation: 54 %, p<0.001, Wilcoxon matched-pairs test, n = 78; Fig. 2). These increments in POm activity seem to represent the bilateral overlapping of simultaneous signals. Similar effects were found stimulating identical (mirror) or different whiskers (nonmirror whiskers) at both sides. These findings were consistently found across animals (n = 8).

**Fig. 2.**
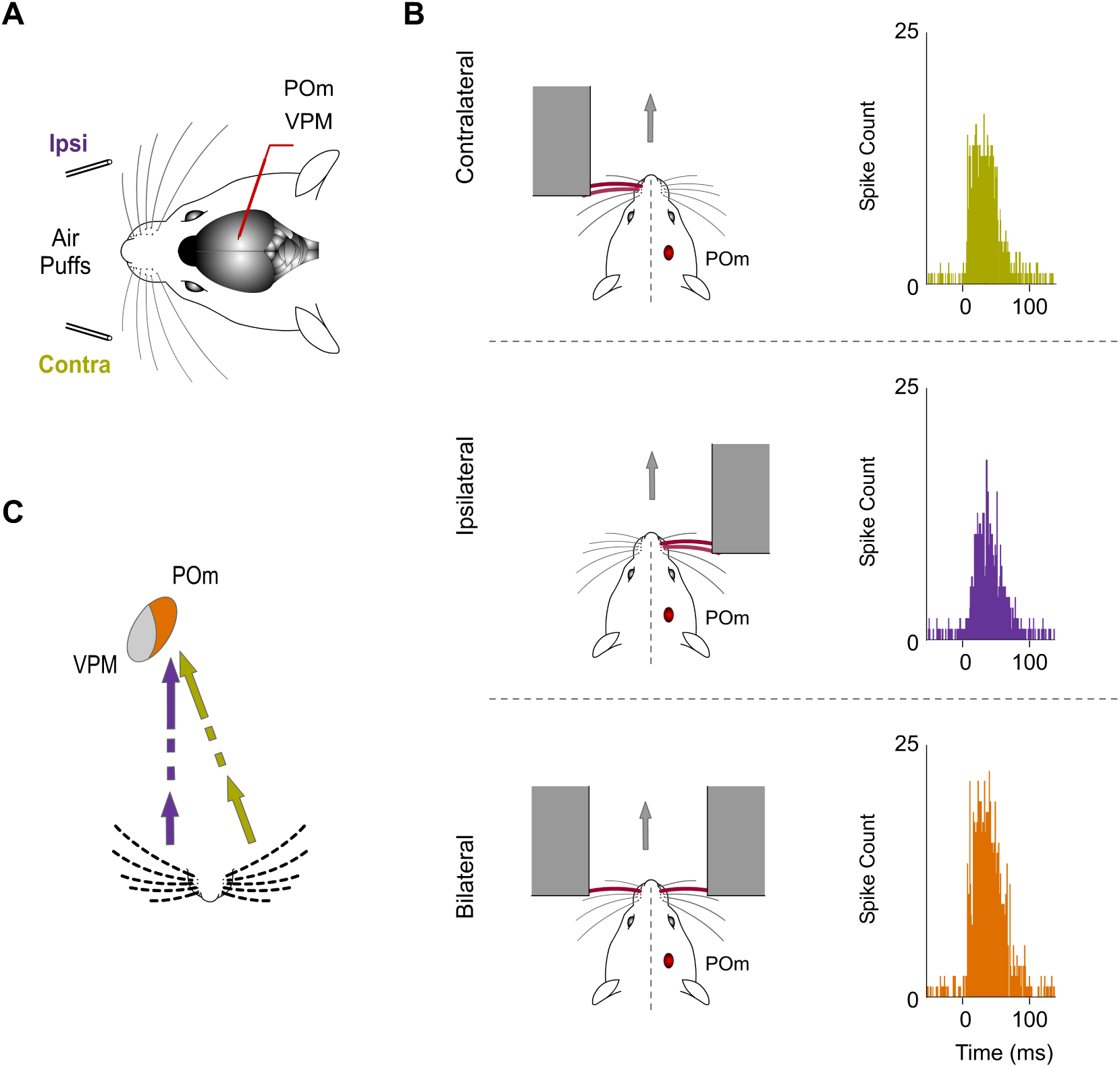
POm integration of bilateral events. (A) Bilateral tactile events were applied via patterns of contra- and ipsilateral simultaneous deflections to investigate whether POm could be implicated in the integration of these signals. (B) PSTHs of representative POm responses evoked by contra-, ipsi- and bilateral whisker stimulations (stimulus duration 40 ms; 30 trials shown for each stimulus) and their corresponding schematic illustrations of their simulated possible real occurrence during the exploration of different objects and apertures in natural conditions are shown. Note the significant increase in firing rate when contra- and ipsilateral whiskers were activated concurrently. (C) These findings indicate that sensory information from both whisker pads is integrated by POm.

In sum, our results show that sensory information from both whisker pads is integrated by POm (Fig. 2C) and demonstrate that POm is implicated in the encoding of bilateral tactile events.

## Discussion

### POm mediates bilateral sensory processing

Although it is well described that POm encodes stimulations of the contralateral whisker pad, our results show that POm is also able to respond to tactile stimulation of ipsilateral whiskers. Since the processing and integration of tactile information from the two sides of the body is central in perceptual function (for example, perceptual tasks requiring bilateral integration), our results show that POm has a fundamental role in bilateral tactile perception.

The difference between ipsi- and contralateral response onset latencies (∼ 10 ms, Fig. 1C, D) suggests that contra- and ipsilateral sensory activities are mediated by different pathways. How ipsilateral information reaches this nucleus remains a question to investigate.

### POm integration of bilateral inputs

Bilateral integration is needed when rodents are exploring tunnels, discerning holes and apertures or detecting their width and shape. Our findings revealed the implication of POm in the representation of bilateral tactile patterns by integrating overlapping information arising from both whisker pads.

The implication of the somatosensory cortex in the processing of bilateral stimuli has been much more studied (Armstrong-James and George 1988; Shuler et al. 2001; Debowska et al. 2011). Moreover, the integration of contra- and ipsilateral whisker inputs has been described previously in the barrel cortex (Shuler et al. 2002). However, the implication of the thalamus in this integration had not been described before. Here, our findings demonstrate the functional implication of the higher-order sensory thalamus in the computation of tactile interactions between body sides. From a neurological perspective, our findings could explain why unilateral thalamic lesions can result in bilateral tactile deficits. This is clinically relevant for the field.

### Differences between VPM and POm

We also characterized VPM thalamic responses delivering the same ipsilateral activation in 6 rats. In contrast to POm, VPM did not respond to ipsilateral stimuli (Fig. 1G). Since the integration of tactile information from the two sides of the body is fundamental in bilateral perception, our results suggest a different implication of these thalamic nuclei in this function.

### POm responses to ipsilateral stimulation are highly sensitive to anaesthesia

It has been reported that during wakefulness POm is strongly more active than during anesthetized state (Masri et al. 2008; Sobolewski et al. 2015). Moreover, under light sedation, POm activity is significantly higher than during general anesthesia (Zhang and Bruno 2019). This evidence indicates that POm is highly sensitive to anaesthesia.

In our experiments, we found that POm responses to ipsilateral stimulation were highly affected by the level of anaesthesia (data not shown). Increasing this level by supplementary doses of urethane abolished POm responses to ipsilateral whisker stimulation. Therefore, high levels of sedation impair the real dynamics of POm functioning. This can explain why these thalamic responses to ipsilateral stimulation had not been described before.

## Materials and Methods

### Ethical Approval

All experimental procedures were carried out under protocols approved by the ethics committee of the Autónoma de Madrid University and the competent Spanish Government agency (PROEX175/16), in accordance with the European Community Council Directive 2010/63/UE.

### Animal procedures, anesthesia and electrophysiology

Experiments were performed on both sexes (5 males and 9 females) adult Sprague Dawley rats (220-300 g). Animals were anesthetized (urethane, 1.3 – 1.5 g/kg i.p.) and placed in a Kopf stereotaxic frame. Local anaesthetic (Lidocaine 1%) was applied to all skin incisions. The skull was exposed and then openings were made to allow electrode penetrations in the thalamus.

Extracellular recordings were made in the posteromedial thalamic complex (POm; P 2.5-4.5, L 2-2.5, D 5-6.5) and in the ventral posteromedial thalamic nucleus (VPM; P 2.8-4.6, L 2-3.5, D 5.5-7; Paxinos and Watson 2007). Tungsten microelectrodes (2–5 MΩ) were driven using an electronically controlled microdrive system. Importantly, recordings of neuronal activity in these brain structures were performed several hours after the application of urethane (typically 4 - 5 h) to obtain a lower level of sedation but in the absence of whisker movements and pinch withdrawal reflexes. The body temperature was maintained at 37°C with a thermostatically controlled heating pad.

### Sensory stimulation

Sensory stimulation was characterized by contra-, ipsi- and bilateral multiwhisker deflections using a pneumatic pressure pump (Picospritzer) that delivers air pulses through polyethylene tubes (1 mm inner diameter; 1-2 kg/cm2). We applied 20-70 trials per stimulus condition at low frequency (0.5 Hz).

### Histology

After the last recording session, animals were deeply anesthetized with sodium-pentobarbital (50 mg/kg i.p.) and then perfused transcardially with saline followed by formaldehyde solution (4%). After perfusion, brains were removed and postfixed. Serial 50 μm-thick coronal sections were cut on a freezing microtome (Leica, Germany). These sections were then prepared for Nissl staining histochemistry for discrimination of thalamic nuclei.

### Data acquisition and analysis

Data were recorded extracellularly from VPM and POm. The raw signal of these *in vivo* extracellular recordings was filtered (0.3–5 kHz band pass), amplified via an AC preamplifier (DAM80; World Precision Instruments, Sarasota, USA) and digitized at 10 kHz. From each recording, we extracted the activity of several clusters of multi-units that were collected by amplitude sorting with the aid of commercial software Spike2 (Cambridge Electronic Design, Cambridge, UK).

We defined response magnitude as the total number of spikes per stimulus occurring between response onset and offset from the peristimulus time histogram (PSTH, bin width 1 ms). Response onset was defined as the first of three consecutive bins displaying significant activity (three times higher than the mean spontaneous activity) after stimulus and response offset as the last bin of the last three consecutive bins displaying significant activity. Response duration was defined as the time elapsed from the onset to offset responses. In all figures, raster plots represent each spike as a dot and each line corresponds to one trial. Spikes were aligned on stimulus presentation (Time 0 ms).

Statistical analysis was performed using GraphPad Prism software (California USA). All data are expressed as the mean ± standard error of the mean (SEM). Error bars in the figures correspond to SEM.

## Additional information

### Data availability statement

The data that support the findings of this study are available from the corresponding author upon reasonable request.

### Author contributions

Carlos Castejon designed and conducted the experiments, analyzed the results and wrote the paper. Angel Nuñez designed and conducted the experiments, analyzed the results, reviewed and edited the paper.

## References

Ahissar E, Sosnik R, Haidarliu S (2000) Transformation from temporal to rate coding in a somatosensory thalamocortical pathway. Nature 406:302–306.

Armstrong-James M, George MJ (1988) Bilateral receptive fields of cells in rat Sm1 cortex. Exp Brain Res 70:155–165.

Castejon C, Barros-Zulaica N, Nuñez A (2016) Control of somatosensory cortical processing by thalamic posterior medial nucleus: a new role of thalamus in cortical function. PLoS One 11:e0148169.

Chiaia NL, Rhoades RW, Bennett-Clarke CA, Fish SE, Killackey HP. (1991) Thalamic processing of vibrissal information in the rat. I. Afferent input to the medial ventral posterior and posterior nuclei. J Comp Neurol. 314:201–216.

Casas-Torremocha, D. et al. (2019) Posterior thalamic nucleus axon terminals have different structure and functional impact in the motor and somatosensory vibrissal cortices. Brain Struct. Funct. 224, 1627–1645.

Debowska, W., Liguz-Lecznar, M. & Kossut, M. (2011) Bilateral plasticity of Vibrissae SII representation induced by classical conditioning in mice. J. Neurosci., 31, 5447–5453.

Diamond ME, Armstrong-James M, Ebner FF (1992) Somatic sensory responses in the rostral sector of the posterior group (POm) and in the ventral posterior medial nucleus (VPM) of the rat thalamus. J Comp Neurol 318:462–476.

Masri, R., Bezdudnaya, T., Trageser, J.C., and Keller, A. (2008). Encoding of stimulus frequency and sensor motion in the posterior medial thalamic nucleus. J. Neurophysiol. 100, 681–689.

Paxinos G, Watson C. (2007) The rat brain in stereotaxic coordinates. San Diego: Academic Press.

Shuler MG, Krupa DJ, Nicolelis MAL (2001) Bilateral integration of whisker information in the primary somatosensory cortex of rats. J Neurosci 21:5251–5261

Shuler MG, Krupa DJ, Nicolelis MAL (2002) Integration of bilateral whisker stimuli in rats: role of the whisker barrel cortices. Cereb Cortex 12:86–97

Sobolewski A, Kublik E, Swiejkowski DA, Kaminski J, Wrobel A. (2015) Alertness opens the effective flow of sensory information through rat thalamic posterior nucleus. Eur J Neurosci. 41:1321–1331.

Veinante P, Jacquin MF, Deschenes M. (2000) Thalamic projections from the whisker-sensitive regions of the spinal trigeminal complex in the rat. J Comp Neurol. 420: 233–43.

Wimmer VC, Bruno RM, de Kock CP, Kuner T, Sakmann B (2010) Dimensions of a projection column and architecture of VPM and POm axons in rat vibrissal cortex. Cereb Cortex 20:2265–2276.

Zhang, W., and Bruno, R. M. (2019). High-order thalamic inputs to primary somatosensory cortex are stronger and longer lasting than cortical inputs. Elife 8:e44158.

